# BrainCellR: A Precise Cell Type Nomenclature R Package for Comparative Analysis Across Brain Single-Cell Datasets

**DOI:** 10.1101/2023.11.08.566209

**Authors:** Yuhao Chi, Simone Marini, Guang-Zhong Wang

## Abstract

Single-cell studies in neuroscience require precise cell type classification and consistent nomenclature that allows for meaningful comparisons across diverse datasets. Current approaches often lack the ability to identify fine-grained cell types and establish standardized annotations at the cluster level, hindering comprehensive understanding of the brain’s cellular composition. To facilitate data integration across multiple models and datasets, we designed BrainCellR. This package provides researchers with a powerful and user-friendly tool for efficient cell type classification and nomination from single-cell transcriptomic data. BrainCellR goes beyond conventional classification approaches by incorporating a standardized nomenclature system for cell types at the cluster level. This feature enables consistent and comparable annotations across different studies, promoting data integration and providing deeper insights into the complex cellular landscape of the brain.

## Introduction

Single-cell RNA sequencing (scRNA-seq) has revolutionized the field of neuroscience by enabling comprehensive characterization of cellular heterogeneity in the brain [1, 2]. This technology provides researchers with unprecedented resolution to study individual cells and uncover cell type-specific gene expression profiles. A standard step in scRNA-seq downstream analysis is cell clustering, i.e., cells with a similar gene expression profile are grouped into clusters; groups of marker genes are then extracted by clusters via statistical analysis, and given a label—typically a cell type or subtype. However, the accurate naming of cell types at the cluster level remains challenging [3-6].

Conventionally, cell type classification in the brain has been based on major classes, such as excitatory neurons or inhibitory neurons, and subclasses, such as Lamp5 or L5 IT [7, 8]. These classifications are determined through morphology, location in the brain, electrophysiological characteristics, and gene expression [9]. However, it is increasingly recognized that these broad categories do not adequately capture the full diversity of cell types within the brain [10]. To achieve a more nuanced understanding of cellular composition, researchers are now focusing on identifying and characterizing fine-grained cell types [11].

A major hurdle in precise cell type classification is the lack of a standardized nomenclature system at the cluster level. Due to the absence of uniform guidelines, different studies often use disparate naming conventions [12, 13]. For example, cell (sub)types might be characterized by their cluster ID numbers [14, 15], grouped by cluster at the subclass level (e.g., Lamp5_1, Lamp5_2), or named using a combination of cell type and marker gene (e.g., L6b P2ry12, or Sst Nts) [7]. This lack of a common nomenclature system leads to inconsistencies and difficulties in comparing cell types across datasets [16, 17], hindering the integration of data from multiple studies and the generation of comprehensive cell atlases.

To address these challenges, we have developed BrainCellR, an R package designed specifically for cell type nomenclature in brain scRNA-seq data. BrainCellR offers a comprehensive set of tools and functionalities to enable researchers to classify and nominate cell types and do comparative analysis across brain single-cell datasets.

## Materials and methods

### The selection of clustering methods

In evaluating clustering methods, we consider both external and internal cluster validity indicators: consistency and the ROGUE score [18]. The external cluster indicator is the consistency of clustering between two datasets, defined as the proportion of corresponding clusters between datasets as shown in Supplementary Figure S1A. The ROGUE score, based on entropy, serves as our internal indicator for clustering as shown in Supplementary Figure S1B [18]. Intuitively, a pure cell cluster is defined as a population with identical function and state across all cells and no variable genes. To select the best clustering approach, we evaluated seven methods and pipelines: the Seurat package using the Louvain algorithm [19], Monocle3 package using Louvain algorithm[20], SC3 using SVM algorithm[21], scCCESS-SIMLR [22], scCCESS-Kmeans [22], scrattch-hicat [10], and our one-iteration method, Consensus1, based on improvements to scrattch-hicat.

In addition to performance metrics, we also considered the execution time as shown in Supplementary Figure S1C and computational memory usage as shown in Supplementary Figure S1D of each method. We thoroughly evaluated these methods across six brain cell datasets as shown in Supplementary Table S1, comprising three human and three mouse datasets. Among the evaluated clustering methods, the ROGUE scores [18] suggest that there may not be a significant difference in performance between most of the methods. However, when considering the external indicators, both Consensus1 and scrattch-hicat exhibit significantly higher scores compared to the other methods. Consequently, these two methods are selected as candidate methods for the clustering process in our pipeline. Scrattch-hicat[10], which shows the best performance in terms of external indicators, is a strong candidate for accurate cell type identification across datasets. On the other hand, Consensus1, with the second-best performance, offers the advantage of significantly reduced computing time compared to scrattch-hicat as shown in Supplementary Figure S1C. Therefore, we have chosen Consensus1 for the clustering process in our pipeline.

#### Evaluating the Consistency of Clusters Between Datasets

The consistency of clustering between datasets is used to evaluate whether the clustering of two data sets is similar after independent clustering. High consistency indicates that cells with similar expression characteristics are grouped into the same class in both sets of data. In this process, we first integrate the two sets of data together for unified clustering (clustering is performed using the methods described in the articles from which each dataset is sourced). Each cell can then obtain a clustering label from this integrated clustering.

Subsequently, we use the clustering algorithm to cluster the two sets of data separately. We then count the distribution of the number of integrated cluster labels contained by cells in each independent cluster and normalize this distribution by dividing it by the maximum value in the distribution. We then calculate the correlation coefficient between the quantity distribution calculated by each cluster from each dataset. If the correlation coefficients between two clusters from different datasets are the highest respectively, then we consider these two clusters to be a pair of clusters with cluster consistency. The proportion of cluster consistency is the ratio between the number of identified clusters with consistency and the total number of clusters contained in the dataset.

### Evaluating the purity of clusters between datasets

The purity score is an entropy-based statistic called ROGUE [18] to quantify the purity of identified cell clusters. The entropy can be defined as

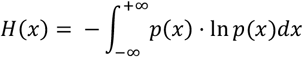

where X is the expression value and p(x) is the probability density function. Then the degree of disorder or randomness of gene expression can be represented as

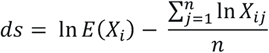

E(*X*_*i*_) is the expectation expression of X under negative binomial distribution for gene i. The ROGUE value which represents the purity score can be defined as

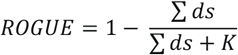

Where K is a parameter to constrain the value between 0 and 1. Additionally, K can also serve as a reference factor to aid in interpreting the purity evaluation.

### Description of the Consensus1 Method

The Consensus1 method is an improvement on the scrattch.hicat [10] method. With the scrattch.hicat method, a subset of cells is selected 100 times for clustering, and then the differentially expressed genes between neighboring clusters are calculated. If no differentially expressed genes can be found between clusters, the two clusters are merged and the process is iterated continuously until no new clusters are generated. If a group of cells is grouped into the same cluster in all 100 iterations, we consider them to be a robust cluster.

The Consensus1 method modifies this process by selecting all the cells for clustering only once, instead of selecting 80% of the cells and clustering them 100 times as in the scrattch.hicat method. The Consensus1 method also preserves the process of merging clusters from the scrattch.hicat method. In this case, the first round of clustering is performed and then the differentially expressed genes between the current cluster and its two adjacent clusters are calculated. If no differentially expressed genes are detected, the cluster is merged with the nearest cluster. This process is iterated until no new cluster is generated.

### The selection of supervised classification methods

The purpose of supervised classification within our pipeline is to assign cell types based on existing annotations for the major classes and subclasses of brain cells. In the case of the cortex, there is a general consensus on the major classes and subclasses of cell types found in specific regions[8, 23]. For instance, in the primary motor cortex of mice, there are two major classes of neurons: Glutamatergic and GABAergic, along with several non-neuronal classes based on the neurotransmitters they release [24]. The classification of these major classes is widely agreed upon in the mouse cortex [10]. Glutamatergic neurons can be further classified into different subclasses based on their cortical localization and projection patterns [7, 25]. Similarly, GABAergic neurons can also be divided into various subclasses. Importantly, these subclasses have been found to be consistent and prevalent across different brain regions of the cortex [10].

We tested our approach using one biological replicate of each dataset (Table S1) as a training set, training the model on the training set using five methods: CHEATH [26], scmap [27], Seurat [19], SingleCellNet [28] and SingleR [29]. We then applied the trained model to another biological replicate of each dataset to evaluate the accuracy of the classification. Based on the evaluation results as shown in Supplementary Figure S2, Seurat [19] performs best in three out of four indicators: Accuracy, Precision, Recall, and F1-score. Several studies have evaluated different methods for cell type classification, with SingleR achieving the highest classification accuracy and Seurat ranking second [30]. While SingleR [29] has shown superior performance, it requires more computational time, particularly for large datasets [30]. By incorporating Seurat and SingleR into our pipeline, we achieve precise and consistent cell type classification across brain single-cell datasets.

### Selection of marker gene identification methods

In BrainCellR, the identification of marker genes involves two substeps: identifying differentially expressed (DE) genes within the same cell subclass; and selecting marker genes from these differentially expressed genes. We evaluated four differential expression gene identification methods: Wilcoxon, ROC curve, T-test, and linear regression, as implemented and described in Seurat [19]. These methods were combined with three marker gene screening methods: 1) Sorting by Specific Score: The top three genes are sorted based on a specific score, which represents their expression specificity across clusters; 2) Top 10% Ranking: Genes sorted by specific score are required to be ranked in the top 10% of all expressed genes; 3) P-value Selection: The top three genes are selected based on the p-value identified by the differential gene detection algorithm. We also explored additional approaches for marker gene detection based on Machine Learning: Random forest [31], PCA [32], and Node2Vec+CNN[33]. Finally, we consider whether to use cells or pseudo-cells as an input. We therefore evaluated two methods based on pseudo-cells: Processing the input data by randomly selecting 10 cells of the same cell type and averaging their expression levels to create pseudo-cells; and the hdWGCNA method [34]. We tested the performance of dozens of methods by combining various input data processing approaches (Raw data, Pseudo-cell, Hdwgcna-generated), four distinct methods for identifying differential gene expression, and multiple marker gene selection methods. We found that the best performance was achieved when using pseudo-cell data as input, identifying differentially expressed genes with ROC, and then selecting the top 10% of highly expressed genes ordered by specific score as shown in Supplementary Figure S3. We incorporate this combination of methods into our pipeline. For cell types where differentially expressed genes could not be identified by the ROC method, we utilized Wilcox test, t-test, and linear regression which are ranked by our evaluation result to identify DE genes and extract marker genes. By incorporating these methods into our pipeline, we ensured to empirically select the most accurate approach for marker genes identification, which is essential for distinguishing and characterizing specific cell types.

### Detailed method for identifying marker gene

The Wilcoxon, ROC, t-, and linear regression tests used to identify differentially expressed genes were the methods provided by Seurat package[19]. Specific score is used to measure the degree of specificity of genes, and the formula is as follows:

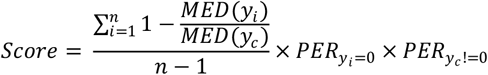

Where, c represents the cell type which currently concerned, i represents other cell types except for c, MED represents the median expression level, and PER represents the percentage meeting the conditions.

The input data for both the random forest and PCA consist of expression matrices. Marker genes are identified based on the feature extraction ability of the model. For the random forest, genes are sorted according to Gini importance [35], while for PCA, genes are sorted based on their ranking on PC1. The input for the Node2Vec+CNN method is the co-expression matrix of expression data between each gene, and the objective of the training is to classify the input genes. The training set comprises the marker genes identified after integrating the two sets of data. Since Node2Vec [33] does not limit the number of co-expressed genes that can be imported, we used the correlation coefficients among all differentially expressed genes for classification.

### Identifying the same cell type

In our pipeline, the name of the cell type is composed of the major class of the cell type, the subclass of the cell type, and the top marker genes of the cell type. If the three marker genes of one cell type overlap with the three marker genes of another cell type within the same subclass, these two cell types are considered to be the same cell type.

## Results

The BrainCellR pipeline can be divided into three steps as shown in Figure 1. The first step involves clustering single-cell data at a fine scale. The second step involves supplying the clusters to a supervised cell type classifier, which outputs major and subclass cell types. The third step is the selection of marker genes from the identified differentially expressed genes. Each step of the pipeline is systematically evaluated to select the most appropriate approach for cell type nomenclature as shown in methods section. The final cell label is then constructed by combining the major class and subclass annotations derived from the classifier with the top three marker genes associated with each cell type.

**Figure 1.**
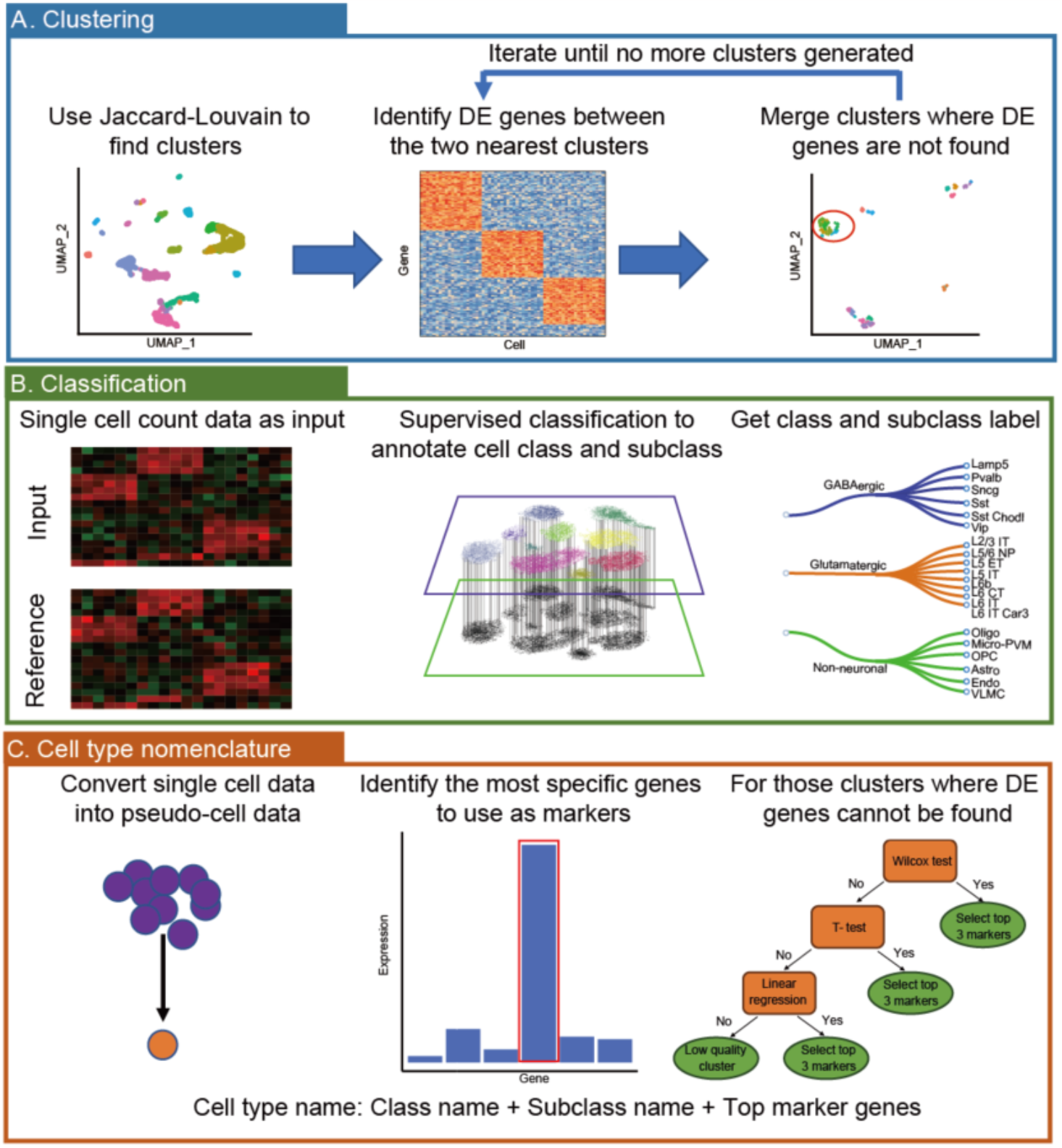
Overall workflow of BrainCellR. The cell type classification and nomination pipeline can be divided into three steps: (A) We use an iterative clustering method called Consensus1 to obtain the cell clustering results; (B) We employ Seurat’s FindTransferAnchors method for supervised classification of the major cell class and cell subclass; (C) After processing single-cell data into pseudo-cells, we identify differentially expressed genes using the ROC methods in Seurat and sequence them according to expression specificity scores. If we cannot find any marker gene for a cell type, we select other differential expression gene identification methods for processing.

The single-cell data types obtained from different biological samples, different individuals, or different sequencing platforms often vary, presenting a challenge in single-cell classification. To evaluate the effectiveness of our pipeline, we conducted studies using (a) six datasets to assess the consistency among different biological data; (b) a mouse dataset to assess the consistency among biological replications; (c) a human dataset to assess the consistency across different individuals; and (d) external mouse datasets, derived from various sequencing methods, to examine the consistency among different sequencing techniques.

### Assessing consistency and comparability of cell types across diverse datasets

We aimed to evaluate whether cell types, labeled identically across different datasets, display similar expression patterns. To determine this similarity, we utilized four metrics: Euclidean distance, Spearman correlation, Pearson correlation, and Cosine similarity, comparing gene expression counts between pairs of cell types. Our analysis incorporated six datasets, which included both human and mouse data as shown in Supplementary Table S1 [1, 2, 10, 36, 37]. In five out of these six datasets, we found that the Euclidean distance between cell types with the same label was significantly smaller than the distance between cell types with different labels. A similar trend was observed for the other three metrics as shown in Figure 2. These results indicate that cell types identified using BrainCellR are consistent and comparable across different datasets, exhibiting significant similarities when labeled identically.

**Figure 2.**
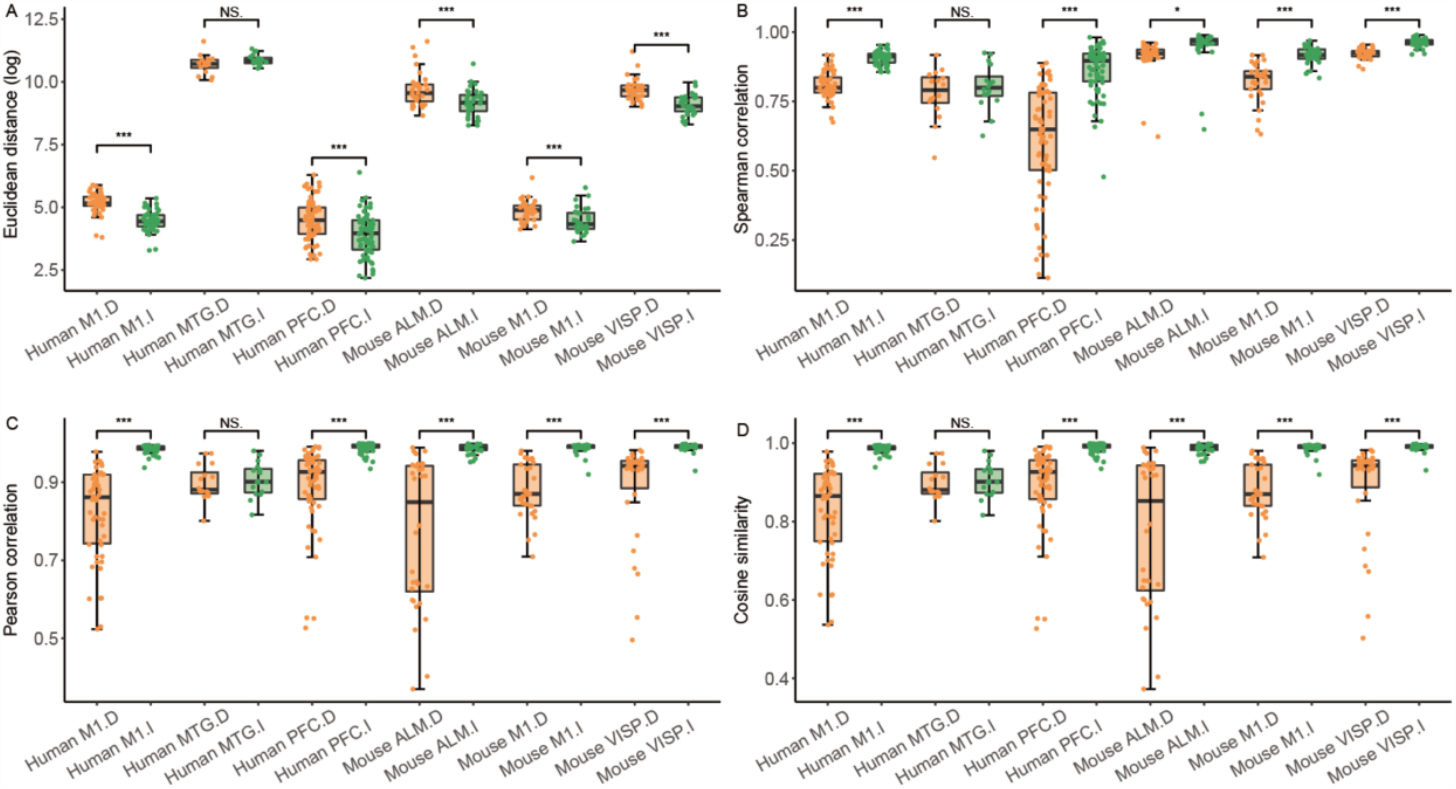
Comparison of similarities across distinct biological datasets. (A) Euclidean distance comparison between identical and different cell types. (B) Spearman correlation comparison between identical and different cell types. (C) Pearson correlation comparison between identical and different cell types. (D) Cosine similarity comparison between identical and different cell types. ‘.I’: identical cell types; ‘.D’: different cell types.

### Evaluating cell type consistency across biological replicate data

We evaluated cell type consistency in biological replication data. Using data from the primary motor cortex of mice as shown in Supplementary Table S2, we selected two sets of biological replicate data, containing 42,108 and 35,303 cells, respectively. We clustered each set independently, then proceeded with the classification of the major cell class and subclass, as well as marker gene selection using the BrainCellR pipeline with the Consensus1 and Seurat methods selected. As a result, we identified 62 cell types in one dataset and 60 cell types in the other, along with their marker genes as shown in Figure 3A. Remarkably, 57 cell types were found to be common between the two datasets, showcasing a high consistency level of 95%. To exemplify this, we selected the Pvalb subclass for display as shown in Supplementary Table S3. Within the data annotated by our pipeline, the Pvalb subclass was further subdivided into seven clusters. Importantly, we observed that the distance between the two datasets for cells belonging to the same cell type was close compared to different cell types in the UMAP plot as shown in Figure 4. This finding indicates that BrainCellR is capable of effectively comparing cell types across different datasets, thus highlighting its utility for cross-dataset analysis.

**Figure 3.**
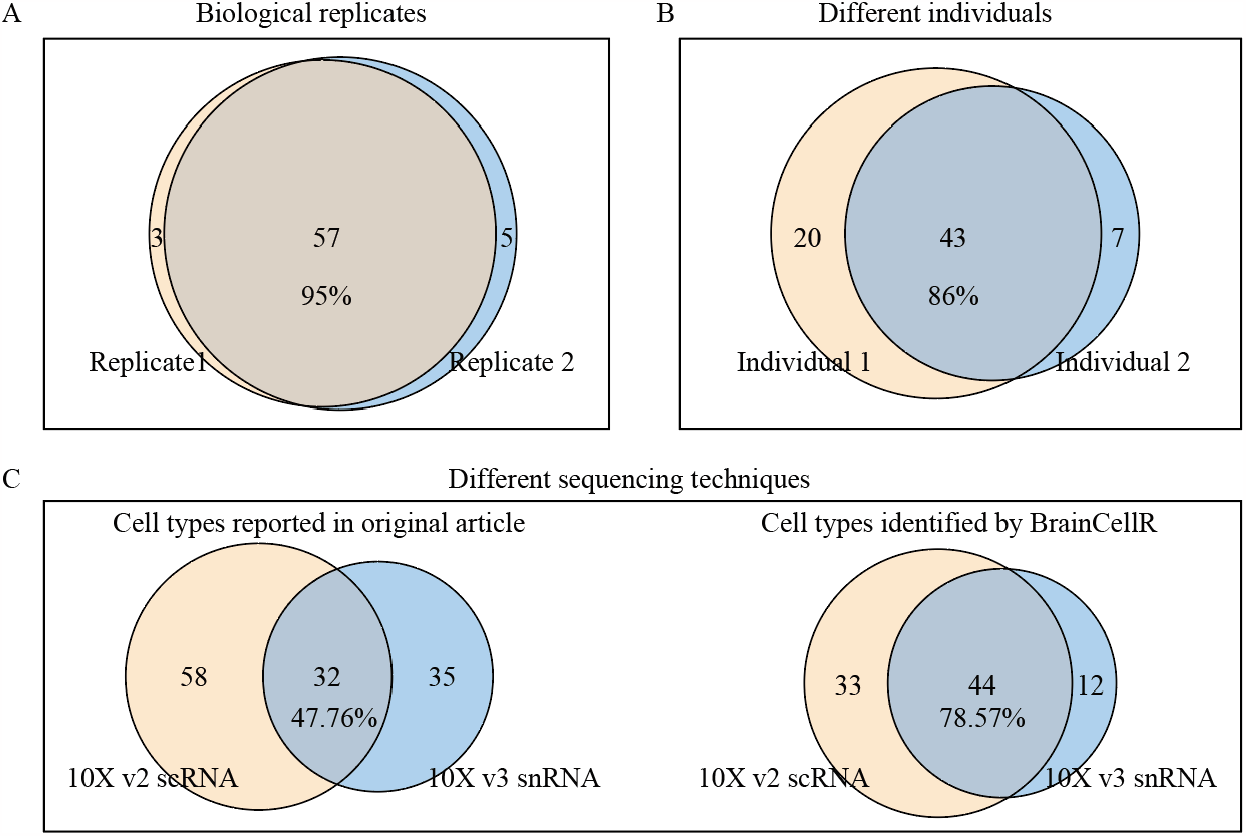
Comparison of cell types from different replicates, individuals, and sequencing techniques. (A) Comparison between different biological replicates, (B) different individuals, and (C) different sequencing techniques.

**Figure 4.**
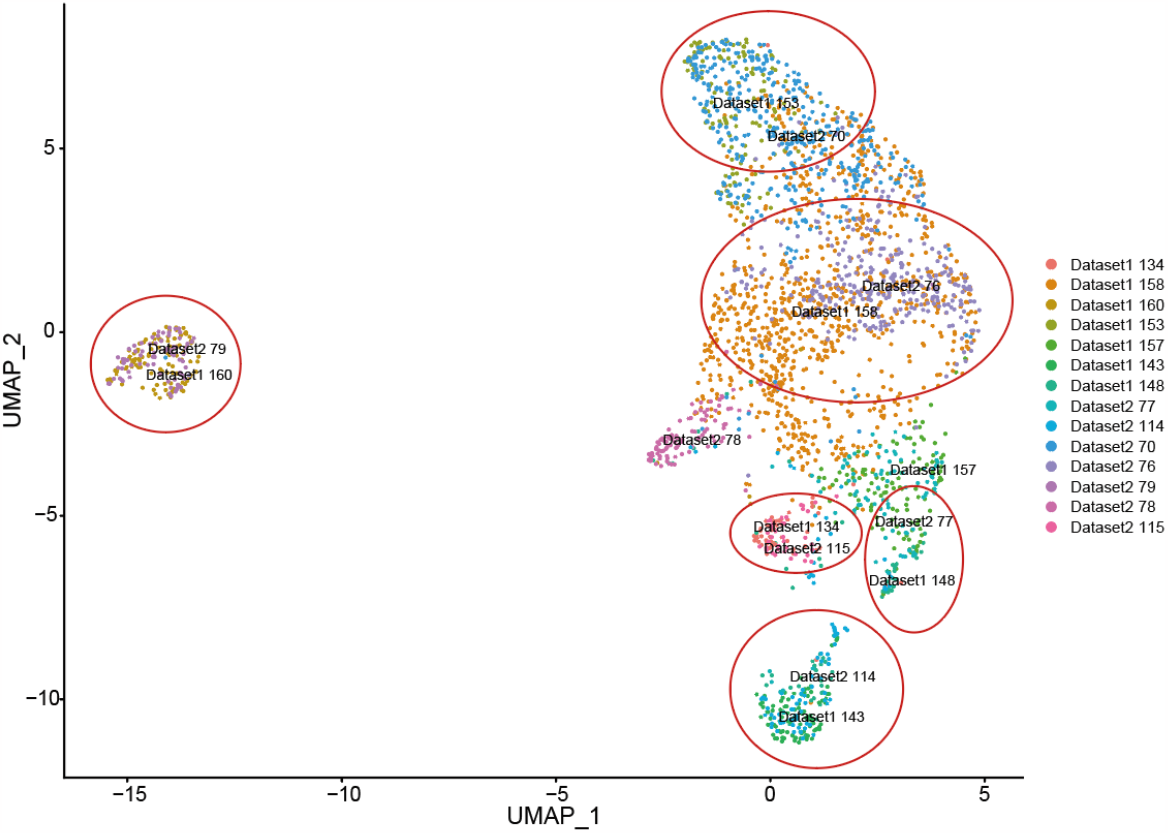
UMAP plot for Pvalb subclass of two combined datasets. The red circles indicate the cell types that correspond between the two datasets, and the cells that are not indicated by the red circles are the cell types that are independent of each of the two datasets.

### Evaluating cell type consistency across different individual sources

Next, we evaluated the cell type consistency in different individual sources. We utilized data from the human middle temporal gyrus (MTG) as our test dataset as shown in Supplementary Table S2, and segregated the cells into two subsets based on the individuals they originated from [37]. These two subsets consisted of 7,206 cells and 7,421 cells, respectively. After conducting marker gene screening using BrainCellR pipeline, we successfully identified a total of 50 and 63 cell types. Remarkably, 43 of these cell types (86%) were found to be consistent across both subsets as shown in Figure 3B.

### Evaluating cell type consistency across different sequencing techniques

Then, we evaluated cell type consistency across data from different sequencing techniques. We conducted a test to examine the consistency of cell type identification within our pipeline across data from high-noise sequencing technology and various batches of experiments [2]. Specifically, we utilized data from 122,641 cells obtained from the primary motor cortex of mice using 10X v2 single-cell sequencing, as well as data from 76,525 cells acquired through 10X v3 single nuclei sequencing as shown in Supplementary Table S2. The analysis of the 10X v2 single-cell data revealed 77 distinct cell types, while the 10X v3 single-nuclei data comprised 56 cell types, with an overlap of 44 cell types (79%). It’s important to note that deviations in cell extraction due to variations in sequencing techniques and cell states, as well as inconsistencies in the number of cells between the datasets, could potentially impact the extraction of certain cell types from the 10X v3 single-nuclei data. In comparison, according to the original data annotations, the two datasets were classified into 90 and 67 categories, with only 32 categories intersecting, resulting in an intersection ratio of merely 47.76% as shown in Figure 3C.

## Discussion

Understanding the diversity of cell types in the brain is a fundamental pursuit in neuroscience research. To facilitate this exploration, we have developed BrainCellR, a powerful pipeline that enables automated classification and nomination of cell types from single-cell transcriptome data. Our pipeline ensures consistent and comparable cell type annotations across different datasets, allowing for comprehensive investigations into the multitude of cell types present in the brain. Please note that the comparison of cell types across datasets is influenced by the number of clusters within the cell subclass. This suggests that our pipeline is better suited for datasets with deep sequencing, a large number of cells, and extensive sampling. BrainCellR holds great promise in advancing our understanding of brain cell diversity and its functional implications. Additionally, BrainCellR has the potential to be extended to other tissues with complex compositions of cell types, not limited to the brain.

## Author contributions

G.W., and S.M. designed the study; G.W., S.M., and Y.C. wrote the manuscript. Y.C. wrote the package and documentation.

